# Lgl cortical dynamics are independent of binding to the Scrib-Dlg complex but require Dlg-dependent restriction of aPKC

**DOI:** 10.1101/867929

**Authors:** Guilherme Ventura, Sofia Moreira, André Barros-Carvalho, Mariana Osswald, Eurico Morais-de-Sá

## Abstract

Apical-basal polarity underpins the formation of specialized epithelial barriers that are critical for metazoan physiology. Although apical-basal polarity is long known to require the basolateral determinants Lethal Giant Larvae (Lgl), Discs Large (Dlg) and Scribble (Scrib), mechanistic understanding of their function is limited. Lgl plays a role as an aPKC inhibitor, but it remains unclear whether Lgl also forms a complex with Dlg or Scrib. Using fluorescence recovery after photobleaching, we show that Lgl does not form immobile complexes at the lateral domain of *Drosophila* follicle cells. Optogenetic depletion of plasma membrane phosphatidylinositol 4,5-biphosphate (PIP_2_) or Dlg removal accelerate Lgl cortical dynamics. However, whereas Lgl turnover relies on PIP_2_ binding, Dlg and Scrib are only required for Lgl localization and dynamic behavior in the presence of aPKC function. Furthermore, light-induced oligomerization of basolateral proteins indicate that Lgl is not part of the Scrib-Dlg complex *in vivo*. Thus, Scrib-Dlg are necessary to repress aPKC activity in the lateral domain but do not provide cortical binding sites for Lgl. Our work therefore highlights that Lgl does not act in a complex but in parallel with Scrib-Dlg to antagonize apical determinants.

## Introduction

Cell polarity is a fundamental property that enables epithelia to act as selective barriers between different environments. Epithelial architecture relies on the asymmetric segregation of evolutionary conserved polarity proteins that define the apical and lateral domains while correctly positioning intercellular junctions (Flores-Benitez and Knust, 2016). Lethal Giant Larvae (Lgl) is one such regulator of apical-basal polarization (Bilder et al., 2000), and is also necessary on other cell polarity contexts, such as asymmetric stem cell division (Ohshiro et al., 2000; Peng et al., 2000), front-rear polarization of migrating cells (Dahan et al., 2014) and anterior-posterior polarization of *Drosophila* oocytes (Fichelson et al., 2010; Tian and Deng, 2008) or *C.elegans* embryos (Beatty et al., 2010; Hoege et al., 2010).

Lgl localizes basally to the adherens junction in most epithelial tissues, where it acts together with Scrib and Dlg to promote lateral identity. These genes were originally described in *Drosophila*, where homozygous mutants display severe epithelial disorganization and neoplastic growth (Bilder et al., 2000; Gateff, 1978; Woods and Bryant, 1989). Further analysis placed these genes together in a basolateral polarity module due to their mutually interdependent localization and antagonistic genetic interactions with the apical Crumbs and aPKC complexes (Bilder et al., 2003; Tanentzapf and Tepass, 2003). Although Scrib, Lgl and Dlg accumulate exclusively at the occluding septate junction in mature *Drosophila* epithelia, such as the adult midgut (Chen et al., 2018), their cortical localization plays evolutionarily conserved roles in apical-basal polarity (Chalmers et al., 2005; Dow et al., 2003; Grifoni et al., 2007; Legouis et al., 2003; McMahon et al., 2001; Musch et al., 2002; Raman et al., 2016; Russ et al., 2012; Sripathy et al., 2011; Yamanaka et al., 2006). Loss of function phenotypes are, however, masked in mammalian epithelia by the presence of multiple paralogues, or, in the case of Scrib, by the compensatory function of proteins with a similar leucine-rich repeat and PDZ protein (LAP) domain (Choi et al., 2019).

The molecular basis for the function of basolateral polarity proteins is still unclear. The predominant hypothesis is that Scrib and Dlg act by ensuring the localization of Lgl at the lateral cortex (Bilder et al., 2000; Kallay et al., 2006; Zeitler et al., 2004). In turn, Lgl prevents the extension of apical determinants (Hutterer et al., 2004), likely by inhibiting the aPKC-Par-6 complex (Atwood and Prehoda, 2009; Betschinger et al. 2003; Yamanaka et al. 2003; Yamanaka et al. 2006), and promoting Crumbs recycling from the lateral cortex (Fletcher et al., 2012). X-ray crystallography provided evidence for a phosphorylation-dependent interaction between the mammalian Lgl2 Basic and Hydrophobic (BH) domain and guanylate kinase domain (GUK) of Dlg4 (Zhu et al. 2014). Moreover, Lgl may bind to the LAP unique region (LUR) domain of Scrib (Kallay et al., 2006), in particular to a small region termed LAPSDa, which is necessary to co-immunoprecipitate Lgl with Scrib in mammalian cell culture (Choi et al., 2019). Nevertheless, it remains unknown whether Scrib and Dlg directly bind Lgl *in vivo* to control its function and localization in the lateral domain.

Given the central role of Lgl, mechanisms that regulate its specific accumulation and dynamic behavior are pivotal for apical-basal polarity. Lgl localization is dynamic and defined by cycles of aPKC and Aurora A phosphorylation and dephosphorylation by Protein Phosphatase 1 (Bell et al., 2015; Betschinger et al., 2003; Carvalho et al., 2015; Moreira et al., 2019). Early studies highlighted that Lgl associates with the actomyosin cortex (Betschinger et al., 2005; Strand et al., 1994), whereas recent work uncovered that Lgl cortical localization is primarily mediated by interactions between its BH domain and plasma membrane phosphoinositides (Bailey and Prehoda, 2015; Dong et al., 2015). These binding partners could therefore aid Scrib and Dlg to form multivalent interactions that regulate binding of Lgl at the lateral cortex, but the contribution of each putative binding component has not been defined.

Here, we characterized Lgl cortical dynamics in the lateral cortex of the *Drosophila* follicular epithelium. We found that Lgl dynamics are dependent on both PIP_2_ and Dlg. However, whereas the first is a critical anchor at the membrane that restricts Lgl cortical mobility, the later regulates Lgl turnover by repressing aPKC ectopic localization. Moreover, light-induced protein clustering suggests that Dlg-Scrib complexes do not contain Lgl, further reinforcing that Lgl acts separately of Scrib-Dlg in antagonizing apical proteins.

## Results and Discussion

### Lgl displays membrane-diffusion behavior and faster dynamics than Dlg and Scrib

To characterize Lgl dynamics in the lateral domain of epithelial cells we performed Fluorescence recovery after photobleaching (FRAP) in post-mitotic stages (stages 7 to 9) of the *Drosophila* follicular epithelium (Figure S1). These stages are suited to dissect the steady-state cortical dynamics of lateral polarity proteins independently of their incorporation in the immobile septate junctions that are formed during late oogenesis (Mahowald, 1972). We first compared the dynamics of Lgl with the core components of the Scrib module using GFP-tagged proteins expressed at endogenous levels. Lgl is significantly more dynamic than Dlg and Scrib (Lgl^t1/2^ ~ 10 s; Dlg^t1/2^ ~ 40 s; Scrib^t1/2^ ~ 60s; Figure 1A, 1B and Movie S1). These differences are likely associated with their distinct binding domains and individual functions (Bonello and Peifer, 2019; Stephens et al., 2018). Moreover, rebleaching a previously bleached region leads to an identical plateau of Lgl fluorescence recovery (Figure 1C). Thus, Lgl does not form a significant subpopulation of immobile complexes at the cortex and plasma membrane.

**Figure 1:**
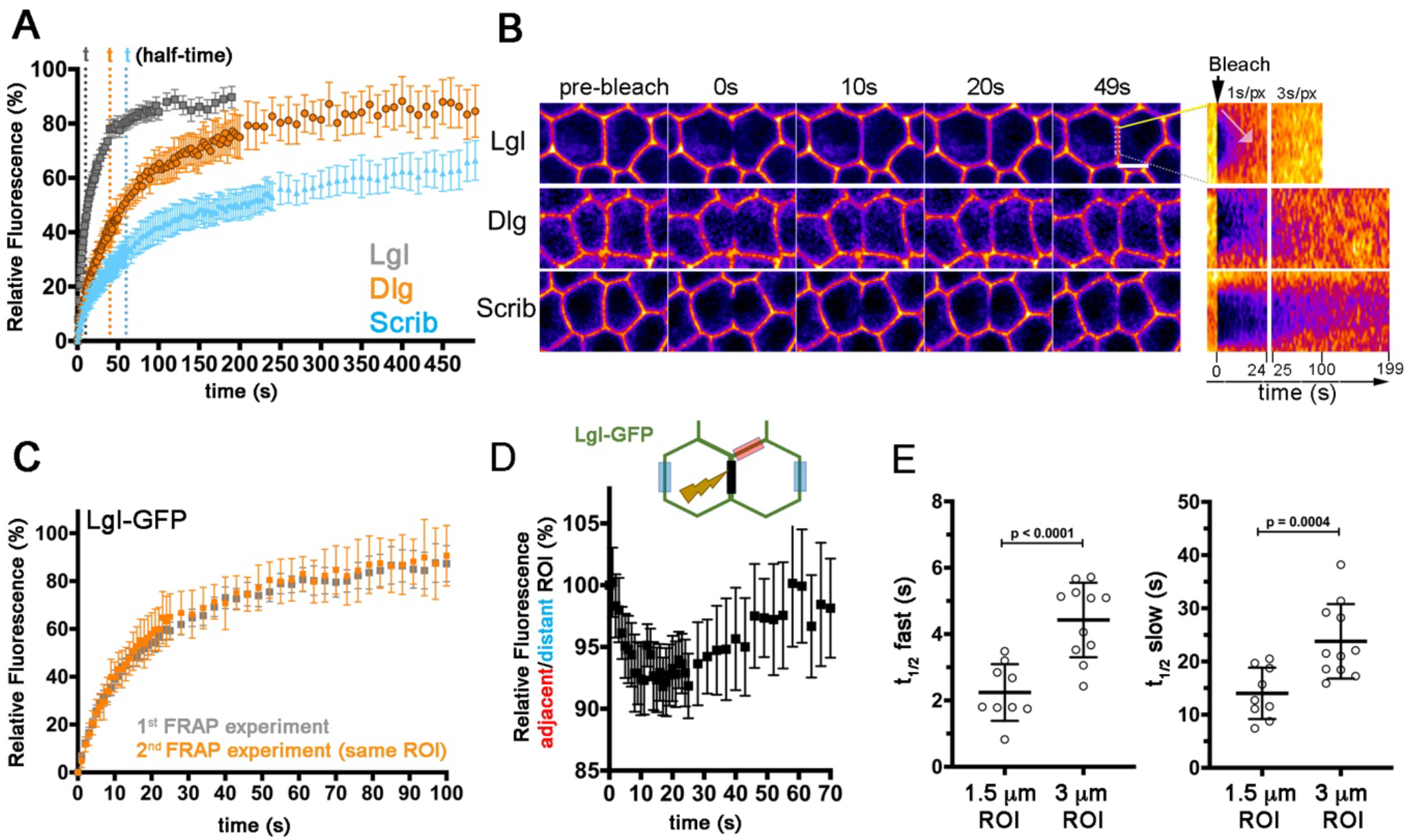
Lgl dynamic behavior is faster than Scrib and Dlg and coupled to membrane-diffusion. **(A)** Normalized mean fluorescence recovery (% ± 95% CI) of endogenously tagged Lgl-GFP (n=43), Dlg-GFP (n=18) and Scrib-GFP (n=22). **(B)** Pseudo-colored images representative of FRAP experiments in the lateral domain of the follicular epithelium show the distinct dynamics of the three basolateral polarity proteins (Movie S1). Kymographs (right) depict the recovery in fluorescence intensity in the photobleached region. Time after photobleaching is indicated in seconds. Scale bar = 5 μm. **(C)** Recovery plots of consecutive FRAP experiments in the same ROI (n=11). **(D)** Plot represents the Lgl-GFP mean intensity (% ± 95% CI) in the interface adjacent (red ROI) to the bleached region normalized to the mean intensity of two distant interfaces (blue ROI; n=23). **(E)** Half-times (t_1/2_) of the fast and slow components of Lgl-GFP dynamics were obtained by a bi-exponential fit of individual FRAP recovery curves of varying photobleaching size (1.5 μm (n=9) vs 3 μm (n=11)). Mean±SD are shown. p-values were calculated using the Mann-Whitney t-test.

FRAP kymographs show that fluorescence recovery initiates at the edges of the photobleached region (Figure 1B - arrows), suggesting that Lgl diffuses along the lateral cortex. Consistent with this, Lgl-GFP fluorescence decreases in neighboring unbleached membranes immediately after photobleaching (Figure 1D). Individual Lgl recovery curves were predominantly best fit by a biexponential equation, suggesting the presence of two components of Lgl dynamic behavior with a fast and slow half-time (Figure 1E). Since diffusion scales with the size of the photobleached region, unlike membrane-cytoplasmic exchange (Fritzsche and Charras, 2015; Sprague and McNally, 2005), we reduced the photobleached region to verify whether biexponential fitting could separate their contributions. However, we concluded that membrane diffusion contributes to both components of Lgl dynamics as both t^1/2fast^ and t^1/2slow^ depend on the size of the photobleached region (Figure 1E). Moreover, we decided to determine a single half-time of fluorescence recovery for subsequent comparative analysis of Lgl dynamics (Table S1) because biexponential fitting was not the best-fit for a significant fraction of the individual Lgl-GFP curves (49% were best-fit by one exponential or produced open-ended 95%-confidence intervals for half-times or plateau after biexponential fitting, n=43).

### Lgl dynamics in the lateral domain are isolated of aPKC and restrained by PIP_2_

The previous results stress the importance of lateral membrane diffusion, but do not exclude a contribution of Lgl exchange between the membrane and cytoplasm. Such turnover could be promoted by aPKC, as aPKC-mediated phosphorylation induces dissociation of Lgl from the apical cortex (Betschinger et al., 2003; Plant et al., 2003; Yamanaka et al., 2003). We applied FRAP analysis to detect residual activity of aPKC at the lateral cortex where, if present, it would promote membrane-cytoplasmic exchange by releasing Lgl from plasma membrane phosphoinositides and cortical interactors. Accordingly, constitutively active *aPKC*^Δ*N*^ (Betschinger et al., 2003) induces cytoplasmic accumulation (Figure 2A) and strongly accelerates the turnover of Lgl-GFP at the lateral cortex (Figure 2B). We then determined whether aPKC-dependent phosphorylation normally plays a role in Lgl turnover in the lateral domain of epithelia by examining the dynamics of Lgl^5A^-GFP, a knock-in allele in which all phosphorylatable serines are mutated to alanine. Lgl^5A^-GFP shows identical kinetics to wild-type Lgl-GFP (Figure 2C). Moreover, although RNAi-mediated depletion of aPKC interferes with apical-basal organization and enables apical accumulation of Lgl, Lgl dynamics are preserved in the lateral cortex (Figure 2C-2E). Thus, even though aPKC inhibits Lgl accumulation in the apical domain, aPKC activity does not impact the dynamics of Lgl at the basolateral cortex in steady-state polarized epithelia.

**Figure 2:**
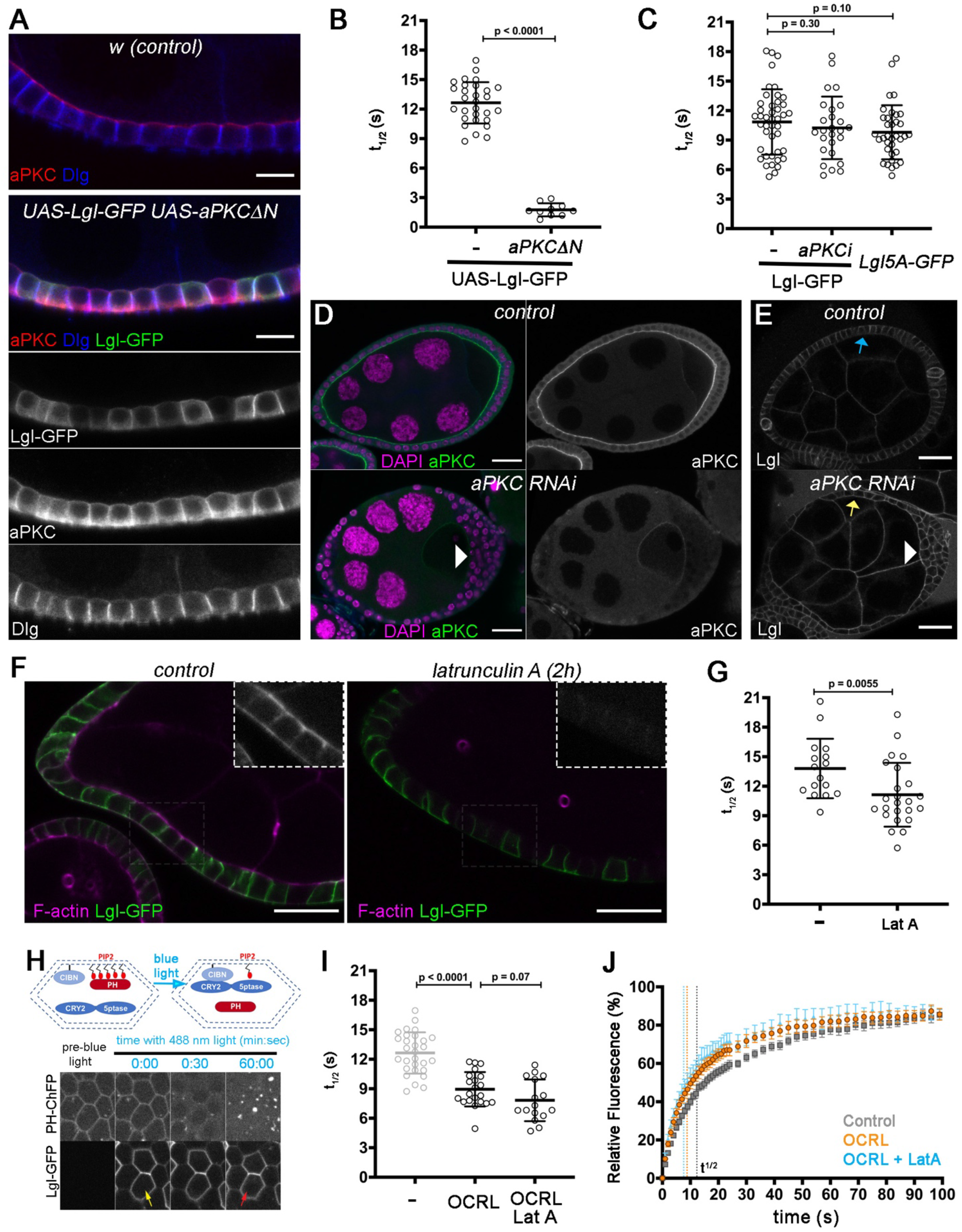
Lgl dynamics are isolated of aPKC activity and restrained by PIP_2_ in the lateral cortex. **(A)** Control or follicle cells expressing UAS-*aPKC*^Δ*N*^ and UAS-Lgl-GFP were stained for aPKC (red) and Dlg (blue). **(B,C)** Scatter plots represent the half-time (t_1/2_) of fluorescence recovery for **(B)** UAS driven Lgl-GFP in control (n= 29) and *aPKC*^Δ*N*^ overexpressing cells (n = 10) and for **(C)** endogenously tagged Lgl-GFP (n = 43) control, aPKC RNAi (n = 26) and Lgl^5A^-GFP (homozygous, n = 35). **(D,E)** Midsagital cross-sections of control and aPKC depleted follicle cells stained for **(D)** DAPI (magenta) and aPKC (green) to validate depletion or **(E)** expressing endogenously Lgl-GFP, which is mislocalized to the apical side upon aPKC depletion (arrow). Arrowhead indicates multilayered tissue. **(F)** UAS-driven Lgl-GFP expressing egg chambers were treated with Latrunculin A (LatA) to disrupt the actin cytoskeleton. F-actin was labelled with phalloidin (magenta and close-up). **(G)** Scatter plots show that Lat A (n = 24) induces a minor, but significant reduction of the Lgl-GFP recovery half-time in comparison to control (n = 16). **(H)** Cartoon depicts the optogenetic depletion of PIP_2_ by light-dependent dimerization between CIBN anchored to the plasma membrane and CRY2 fused with the inositol polyphosphate 5-phosphatase OCRL. Blue-light exposure leads to rapid release of PH-ChFP from the membrane, a slight accumulation of Lgl in the cytoplasm (arrows, movie S2) **(I,J)** and faster cortical dynamics of Lgl. **(I)** Scatter plots show the half-time distribution and **(J)** graph showing the relative mean fluorescence recovery (± 95% CI) of UAS-driven Lgl-GFP in control (same dataset as (B)), upon PIP_2_ depletion (OCRL, n= 21) and also with simultaneous disruption of the actin cytoskeleton (OCRL_LatA, n= 16). Dashed lines indicate the half-times (t_1/2_). Mean±SD are shown in (B,C,G,I). p-values were calculated using the Mann-Whitney t-test. Scale bars = 10 μm (A); 25 μm (D,E,F)

Interactions between Lgl and its putative binding partners should restrict Lgl mobility in the lateral cortex and so we used FRAP to determine the relative contributions of two known Lgl interactors, plasma membrane phosphoinositides and the actin cortex. Early work suggested that Lgl is removed from the cortex in embryonic neuroblasts in response to actin disruption with Latrunculin A (LatA) (Betschinger et al., 2005), but not in the embryonic epithelium (Dong et al., 2015). Disruption of the actin cortex in the follicular epithelium also does not affect the basolateral localization of either Lgl, Dlg or Scrib, suggesting that apical-basal polarization is largely maintained upon acute disruption of the actin cytoskeleton (Figure 2F and S2). However, LatA induced a minor but significant acceleration of the fluorescence recovery of Lgl-GFP (Figure 2G). Thus, the actin cytoskeleton is not required for Lgl localization to the lateral cortex, but restrains Lgl mobility, possibly through binding of Lgl to myosin and other actin cortex associated proteins (Dahan et al., 2014; Strand et al., 1994).

Lgl localization to the plasma membrane depends on the interaction with negatively charged phosphoinositides (Bailey and Prehoda, 2015; Dong et al., 2015; Visco et al., 2016), such as phosphatidylinositol-4,5-biphosphate (PIP_2_), whose depletion is sufficient to induce partial mislocalization of Lgl to the cytoplasm of follicle cells (Dong et al., 2015). We used an optogenetic tool that depletes PIP_2_ with high temporal control through the light-induced recruitment of the 5-phosphatase OCRL to the plasma membrane, where it converts PI(4,5)P_2_ to PI(4)P (Guglielmi et al., 2015). This approach triggers efficient removal of PIP_2_ sensor PH-ChFP from the plasma membrane in the follicular epithelium, while maintaining cortically localized UAS-driven Lgl-GFP (Figure 2H). PIP_2_ depletion increases Lgl turnover, supporting the idea that PIP_2_ is a major binding partner promoting its localization and restraining its turnover at the cortex (Figure 2I and 2J). Consistent with this, disruption of the actin cytoskeleton does not significantly increase the mobility of Lgl-GFP when PIP_2_ levels are simultaneously depleted (Figure 2I and 2J).

### Dlg/Scrib regulate Lgl dynamics and localization by preventing aPKC ectopic activity

It has long been known that Dlg is required for Scrib and Lgl cortical localization in the *Drosophila* embryonic epithelium (Bilder et al., 2000). Their cortical localization is also disrupted in *dlg* mutant follicle cells (Figures 3A, 3B and Figure S3A), but it is unclear how Dlg and Scrib promote Lgl cortical localization. Dlg does not interfere directly with the plasma membrane binding sites, as GFP-tagged PH remains unchanged in *dlg* mutant follicle cells (Figure 3C). Alternatively, mislocalization of aPKC activity in *dlg* mutant cells ((Bilder et al., 2000; Bilder and Perrimon, 2000; Franz and Riechmann, 2010), Figure 3D) could displace Lgl from the plasma membrane. Accordingly, and unlike Lgl-GFP, an aPKC-insensitive form of Lgl, Lgl^5A^-GFP, localizes in the cortex of *dlg* mutant clones (Figures 3E, 3F, S3C and S3D). This suggests that Dlg is not required for Lgl cortical recruitment in the absence of aPKC phosphorylation. To confirm this hypothesis, we generated *apkc, dlg* double mutant cells and observed that aPKC disruption restores Lgl to the cortex (Figure 3G). Long-term disruption of aPKC activity enables constitutive binding of unphosphorylated Lgl at the plasma membrane, which could mask a role for Dlg in Lgl cortical recruitment. We therefore inactivated aPKC acutely using a recent analogue-sensitive aPKC allele (Hannaford et al., 2019). Live imaging of endogenously expressed Lgl-GFP shows that aPKC inhibition induces a quick reallocation of Lgl from the cytoplasm to the cortex in *dlg* mutant cells (Figure 3H).

**Figure 3:**
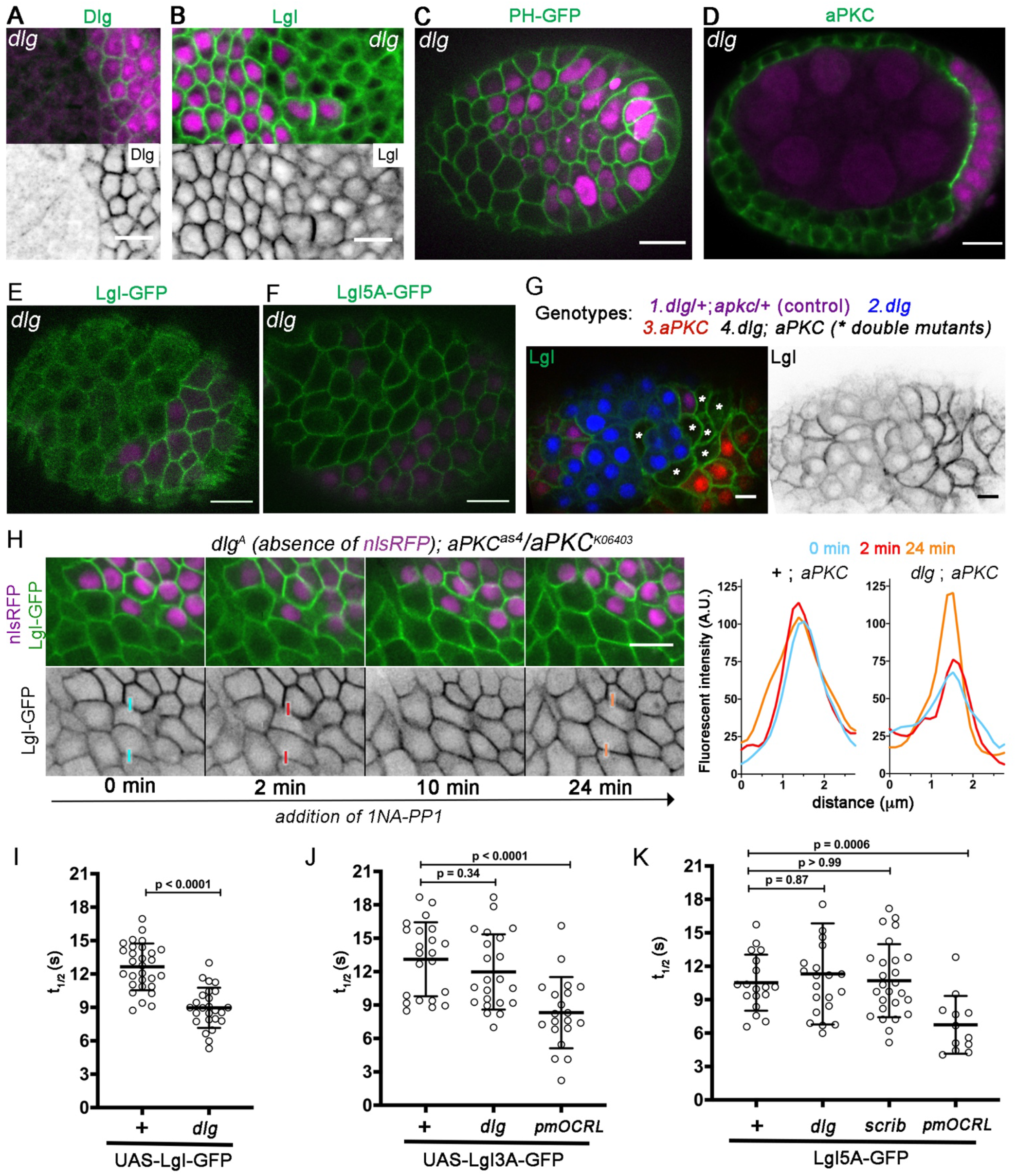
Dlg/Scrib control Lgl localization and dynamics by preventing ectopic aPKC activity. **(A-F)** Follicular epithelium with *dlg^A^* mutant clones (absence of nls-RFP (magenta)) stained for **(A)** Dlg **(B)** Lgl or **(D)** aPKC, or expressing **(C)** PIP_2_ sensor PLCδPH-GFP or **(E)** Lgl-GFP or **(F)** Lgl^S5A^-GFP at endogenous levels. **(G)** Clonal analysis in the follicular epithelium shows that Lgl cortical localization is disrupted in *dlg^A^* mutants (blue - absence of nlsRFP), but is rescued when aPKC is simultaneously removed (*dlg^A^, aPKC^K06403^* (*, black)). Control (purple - *positive for nlsRFP (red) and nlsGFP (blue)*) and *aPKC^K06430^* (red - absence of nlsGFP) mutant cells are present. **(H)** Selected frames of time-lapse showing cells mutant for the ATP analogue-sensitive allele of aPKC (*aPKC^as4^/aPKC^K06403^*) and containing *dlg^A^* clones that were treated with 1NA-PP1. Pixel intensity plots show Lgl-GFP fluorescence (A.U.) at the color-coded time-points across the indicated interfaces of wild-type ((nlsRFP positive) +;*aPKC^as4^*) and mutant (*dlg;aPKC*) cells. **(I, J, K)** Scatter plots of recovery halftime (t_1/2_) show that **(I)** UAS-driven Lgl-GFP cortical dynamics are accelerated in *dlg^A^* cells (n= 25) in comparison to control (n = 29). **(J)** UAS-driven Lgl^3A^-GFP dynamics are unchanged in *dlg^A^* mutants (n= 22) in relation to control (n= 22), and accelerated by optogenetic depletion of PIP_2_ (n= 20). **(K)** Lgl^5A^-GFP cortical dynamics are also not affected in dlg^A^ (n=21) and scrib^2^ (n=26) mutant follicle cells in relation to control (n= 18), but are accelerated upon PIP_2_ depletion (n = 12). Mean±SD are shown. Scale bars = 10 μm. p-values were calculated using the Mann-Whitney t-test.

Altogether, the previous results show that Dlg binding is dispensable for Lgl cortical recruitment, but it could nevertheless regulate Lgl turnover at the cortex. We used FRAP as a sensitive assay to further test the putative role of Dlg as a binding partner of Lgl. We resorted to an UAS-driven Lgl-GFP to monitor Lgl cortical dynamics as it largely restores Lgl cortical localization in *dlg* mutants (Figure S3E). This is consistent with an inhibitory role of Lgl overexpression on aPKC activity and supports the notion that aPKC activity can be titrated by overexpression of its substrates, which would compete for the catalytic site (Holly and Prehoda, 2019). Nevertheless, UAS-driven Lgl-GFP displays faster fluorescence recovery in the absence of Dlg (Figures 3I). Given that Dlg is necessary to localize Scrib, which is also required for Lgl localization (Figure S3F), this result could be compatible with Lgl binding to Dlg or Scrib. To determine the specific contribution of Dlg and Scrib as Lgl cortical binding sites and simultaneously untangle this from their impact on the extension of aPKC activity, we measured the dynamics of aPKC-insensitive versions of Lgl in both *dlg* and *scrib* mutants. The dynamic behavior of nonphosphorylatable Lgl is not significantly different in either *dlg* or *scrib* mutants (Figure 3J and 3K). In contrast, optogenetic depletion of PIP_2_ increases the turnover of nonphosphorylatable Lgl (Figure 3J and 3K). We therefore conclude that whereas PIP_2_ is a major binding site for Lgl at the plasma membrane, Lgl cortical dynamics are independent of direct protein-protein interaction with the Scrib-Dlg module.

### The Dlg-Scrib complex does not contain Lgl in the follicular epithelium

Despite the widespread notion that Dlg, Lgl and Scrib act together as a basolateral protein complex, there are no clear *in vivo* evidence for direct interactions. We have recently repurposed light-induced protein clustering by LARIAT (light-activated reversible inhibition by assembled trap) (Lee et al., 2014; Qin et al., 2017) to probe for protein-protein interactions in S2 cells (Osswald et al., 2019). By combining the light-sensitive cryptochrome CRY2 tagged with a VHH-GFP nanobody and GFP knock-in lines for Lgl, Dlg and Scrib, we now implemented an approach to probe for interactions between basolateral proteins in the follicular epithelium by evaluating co-recruitment to multimeric clusters (Figure 4A). Light stimulation induced the formation of Scrib-GFP clusters, which accumulate below the adherens junction (Figure 4B) and contain Dlg (Figure 4C).

**Figure 4:**
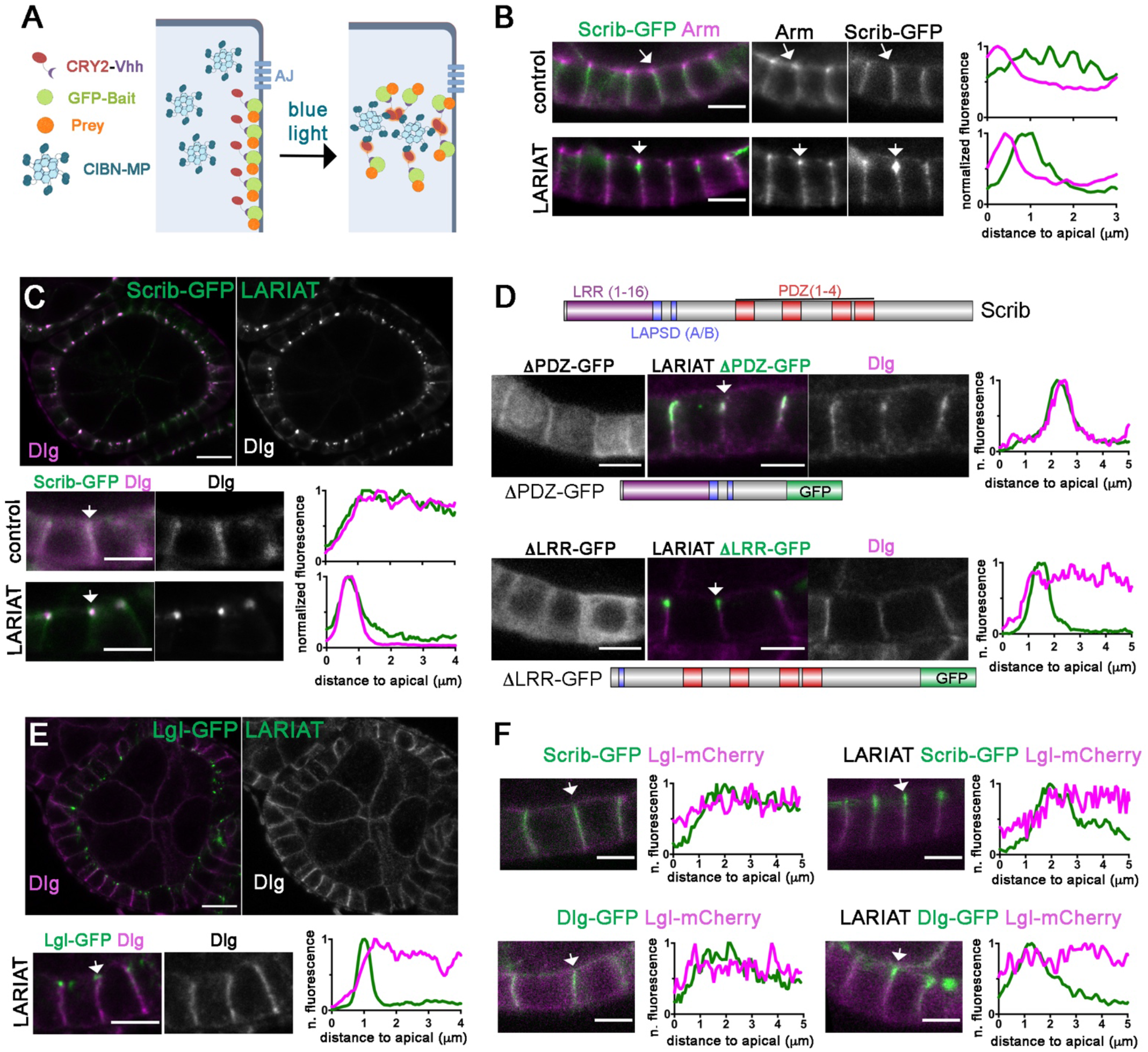
*in vivo* detection of Scrib-Dlg complexes that lack Lgl. **(A)** Detection of protein-protein interactions through light-induced clustering by LARIAT. The GFP-tagged protein (bait) binds CRY2-V_H_H, which homodimerizes and interacts with CIBN-MP upon light-exposure to generate large multimeric complexes that trap interactors (prey) **(B)** LARIAT-mediated protein clusters of Scribble-GFP localize below adherens junctions stained with Armadillo. **(C-E)** Immunofluorescence of Dlg reveals co-aggregation with (C) Scribble-GFP and **(D)** ScribΔPDZ-GFP, but not with Scrib-ΔLRR-GFP or (E) Lgl-GFP. **(F)** Scribble-GFP and Dlg-GFP clusters do not coaggregate with Lgl-mCherry. **(B-F)** Pixel intensity profiles were measured along the cortex (arrows) and were normalized to maximum intensity. Scale bars = 5μm (all close-ups); 10μm (egg chambers in C and E).

To identify how Dlg binds Scrib, we induced clustering of Scrib versions lacking major protein interacting domains, namely the leucine-rich repeats (ΔLRR-GFP) or the PDZ domains (ΔPDZ-GFP), which bind the Dlg binding protein GUK-holder (Caria et al., 2018), and were recently shown to play specific roles in the establishment of polarity, tricellular junction formation and mitotic spindle orientation in *Drosophila* (Bonello et al., 2019; Sharifkhodaei et al., 2019; Nakajima et al., 2019). Both proteins display largely cytoplasmic localization (Figure 4D), but removal of the PDZ domains still enables cortical enrichment as previously reported (Albertson et al., 2004; Zeitler et al., 2004). More importantly, Dlg is absent from ΔLRR-GFP clusters, but co-clusters with ΔPDZ-GFP (Figure 4D). Thus, the Scrib-Dlg complex identified in the follicular epithelium requires interactions with Scrib LRR-LAPSD domains, which are both essential and partially sufficient for apical-basal polarization and proliferation control (Choi et al., 2019; Zeitler et al., 2004).

We next tested whether Lgl is also included in Dlg-Scrib complexes. In contrast to Scrib-GFP clusters (Figure 4C), Dlg is absent from Lgl-GFP clusters (Figure 4E). Conversely, Lgl-mCherry does not co-localize with either Scrib-GFP or Dlg-GFP clusters upon light-exposure (Figure 4F). These experiments further suggest Lgl is not part of the protein complex formed between Scrib and Dlg in the lateral domain of follicle cells.

### Conclusion

Understanding how basolateral proteins maintain apical-basal polarity requires approaches to explore their dynamic behavior, which is intimately connected to their function. Dlg and Scrib mobility has been previously studied in the context of septate junctions (Babatz et al., 2018; Oshima and Fehon, 2011). We now compared the dynamics of the basolateral determinants in the lateral domain and dissected the mechanisms of Lgl association to the cortex. Lgl dynamics are tightly linked to membrane diffusion, as previously observed for PLCδPH-GFP, which binds PIP_2_ with high affinity (Goehring et al., 2010; Hammond et al., 2009). We show that plasma membrane PIP_2_ are indeed major binding sites for Lgl as previously suggested (Bailey and Prehoda, 2015; Dong et al., 2015). Even though regulation of Lgl affinity to PIP_2_ seems to be the critical factor controlling Lgl cortical dynamics, we show that aPKC disruption does not alter Lgl dynamics in the lateral cortex. This suggests the presence of a physical or biochemical barrier that segregates apical and basolateral membranes in epithelial cells. Biochemical competition may also be at work as the isolation of Lgl dynamic behaviour at the lateral cortex of epithelia resembles the maintenance phase of PAR polarity in the *C.elegans* zygote, where the dynamic behavior of PAR proteins is only altered when entering the region enriched in the opposing PAR species (Goehring et al., 2011).

The aforementioned findings underline the significance of mechanisms that efficiently restrain aPKC localization to the apical domain. However, the ability of Lgl to act as an inhibitor and substrate of aPKC has led to a causality dilemma in understanding how Lgl functions together with other apical antagonists. Even if our work cannot exclude the formation of short-lived protein complexes that represent a minor contribution to Lgl dynamics, it shows that Dlg-Scrib are neither critical recruitment-factors nor major binding partners for Lgl in the lateral cortex. Instead they regulate cortical Lgl by preventing the ectopic localization and, ultimately, the activity of aPKC in the lateral domain. Further work is required to characterize how Dlg and Scrib suppress apical determination. Since Dlg is itself an aPKC substrate (Golub et al., 2017), Dlg could buffer aPKC activity towards other substrates. This could promote Lgl accumulation at the cortex, and concomitantly prevent aPKC lateral extension by inhibiting the aPKC-mediated stabilization of apical complexes in the lateral cortex (Fletcher et al., 2012). Moreover, Scrib-Dlg determine the assembly and location of adherens junctions during apical-basal polarization of the embryonic blastoderm (Bonello et al., 2019), which could participate of apical proteins. In conclusion, this study provides evidence that Lgl does not act as part of the Scrib-Dlg complex, suggesting individual parallel mechanisms to antagonize apical identity.

## Materials and Methods

### *Drosophila* stocks and genetics

*Dlg^CC01936^-GFP, Scrib^CA07683^-GFP (Buszczak et al., 2007), Lgl-GFP*, *Lgl^S5A^-GFP* and Lgl-mCherry (Dong et al., 2015) were used for endogenous expression of tagged polarity proteins; *UAS-Lgl^3A^-GFP* and *UAS-Lgl-GFP (Betschinger et al., 2003*) for overexpression conditions; *UAS-pmCIBN* and *UAS-CRY2-OCRL* for optogenetic depletion of PIP_2_ in the plasma membrane ((Guglielmi et al., 2015), kindly provided by Stephano de Renzis); UAS-LARIAT (VhH-SNAPtag-CRY2(PHR)-P2A-CIB1MP) for light-induced clustering of GFP-tagged proteins ((Qin et al., 2017) kindly provided by Xiaobo Wang), including *UAS-Scrib*Δ*LRR-GFP* (BDSC: #59084) and *UAS-Scrib*Δ*PDZ-GFP* (BDSC: #59083); *ubi-PLCγPH-eGFP or ubi-PLCγPH-mCherry* (PH-ChFP) (Herszterg et al., 2013, kindly provided by Yohanns Bellaiche) was used to label PIP_2_ with the Pleckstrin Homology (PH) domain of PLCγ; *UAS-aPKC RNAi* (BDSC: #25946); *UAS aPKC*^Δ*N*^ (Betschinger et al., 2003); *aPKC^K06403^* (Wodarz et al., 2000); *apkc^as4^*, an analogue-sensitive allele used for acute inhibition of aPKC (Hannaford et al., 2019, kindly provided by Jens Januschke); *scrib^2^* (Bilder and Perrimon, 2000). We used a recently generated *dlg^1A^* mutant allele, which was produced with low concentrations of ethyl methylsulphonate to diminish secondary alterations (Haelterman et al., 2014; Yamamoto et al., 2014). *dlg^1A^* encodes a stop codon on the third PDZ domain of Dlg and behaves as a protein null allele. Mitotic clones were generated with the FLP/FRT-mediated mitotic recombination system and were induced by heat shock at 37°C (Xu and Rubin, 1993).

*Drosophila melanogaster* stocks were reared on standard cornmeal/agar/molasses/yeast media at 18°C or 25°C. To boost UAS-RNAi or UAS-LARIAT expression, flies were raised at 29°C ON. Flies were protected from the light before light-induction in the optogenetic experiments. *tj-GAL4* and *GR1-GAL4* were used to drive the expression of UAS transgenes in the follicular epithelium. The list of genotypes used is shown below.

### Genotypes

FRAP analysis of UAS-Lgl-GFP:

- *tj-GAL4/*+; *UAS-Lgl-GFP/*+ (Figures 2B, 2I, 3I)
- *tj-GAL4/*+; UAS aPKC^ΔN^/*UAS-Lgl-GFP* (Figure 2A, B)

*Lgl^5A^* homozygous

- *hs-FLP; Lgl^S5A^-GFP FRT40A/ nls-RFP FRT40A* (Figure 2C)

Lgl-GFP in aPKC RNAi

- *tj-GAL4/*+; *UAS aPKC RNAi/Lgl-GFP* (Figure 2C, 2E)

aPKC RNAi (Figure 2D)

- *tj-GAL4/*+; *UAS aPKC RNAi/* +

Optogenetic PIP_2_ depletion:

- UAS-*pmCIBN/UAS-CRY2-OCRL;tj-GAL4/*+; *ubi-PH-mCherry/*+; *UAS-Lgl-GFP/*+ (Figures 2 H,I,J)
- UAS-*pmCIBN/UAS-CRY2-OCRL;tj-GAL4/*+; *ubi-PH-mCherry/*+; *UAS-Lgl^3A^-GFP/*+ (Figure 3J)
- UAS-*pmCIBN/UAS-CRY2-OCRL;tj-GAL4/*+; *Lgl^5A^-GFP/ ubi-PH-mCherry* (Figure 3K)

*dlg* mutant clones:

- *dlg^A^ FRT19A/hs-FLP nls-RFP FRT19A* (Figure 3 A,B,D)
- *dlg^A^ FRT19A/hs-FLP nls-RFP FRT19A;; ubi-PH-mCherry* (Figure 3C)
- *dlg^A^ FRT19A/hs-FLP nls-RFP FRT19A; tj-GAL4/*+; *UAS-Lgl-GFP/*+ (Figure 3I and S3E)
- *dlg^A^ FRT19A/hs-FLP nls-RFP FRT19A; tj-GAL4/*+; *UAS-Lgl^3A^-GFP/*+ (Figure 3J)
- *dlg^A^ FRT19A/hs-FLP nls-RFP FRT19A; Lgl^5A^-GFP/*+ (Figure 3F, K, S3D)
- *dlg^A^ FRT19A/hs-FLP nls-RFP FRT19A; Lgl-GFP* (Figure 3E, S3C)
- *dlg^A^ FRT19/hs-FLP nls-RFP FRT19A;; Scrib-GFP* (Figures S3A,S3B)

*double dlg,aPKC* mutants

- *dlg^A^ FRT19A/hs-FLP nls-RFP FRT19A; FRT42B aPKC^K06430^ /FRT42B nls-GFP* (Figure 3G)
- *dlg^A^ FRT19/hs-FLP nls-RFP FRT19A; Lgl-GFP aPKC^K06403^/aPKC^as4^* (Figure 3H)

*scrib* mutant clones:

- *hs-FLP/*+; *Lgl^5A^-GFP/*+; *FRT82 RFP/FRT82 scrib^2^* (Figure 3K)
- *hs-FLP/*+; *Lgl-GFP/*+; *FRT82 RFP/FRT82 scrib^2^* (Figure S2F)

LARIAT optogenetic clustering experiments:

- *tj-GAL4 or UAS-LARIAT/Cyo*; *Scrib-GFP/TM6* (control) (Figure 4B)
- *tj-GAL4/UAS-LARIAT; Scrib-GFP/TM6* (Figure 4B)
- *tj-GAL4 or UAS-LARIAT/Cyo*; *Scrib-GFP/ Scrib-GFP* (control) (Figure 4C)
- *tj-GAL4/UAS-LARIAT; Scrib-GFP/ Scrib-GFP* (Figure 4C)
- *UAS-Scrib*Δ*PDZ-GFP/Cyo; UAS-LARIAT GR1-GAL4/*+ (Figure 4D)
- *UAS-Scrib*Δ*LRR-GFP/UAS-LARIAT GR1-GAL4* (Figure 4D)
- *tj-GAL4/Lgl-GFP; UAS-LARIAT GR1-GAL4/*+ (Figure 4E)
- *Lgl-mCherry/Cyo; UAS-LARIAT/Scrib-GFP* (control) (Figure 4F)
- *tj-GAL4/Lgl-mCherry; UAS-LARIAT/ Scrib-GFP* (Figure 4F)
- *Dlg-GFP/*+; *tj-GAL4/Lgl-mCherry; TM6/* + (Figure 4F)
- *Dlg-GFP/*+; *tj-GAL4/Lgl-mCherry; UAS-LARIAT/* + (Figure 4F)

### Immunofluorescence

Ovaries of well-fed *Drosophila* females were fixed in 4% paraformaldehyde (in PBS) for 20 minutes, washed 3 x 10 minutes in PBT (PBS with 0.05% of Tween 20), blocked with PBT-10 (PBT supplemented with 10% BSA) and then incubated overnight with primary antibodies in PBT supplemented with 1% BSA. After 4 x 30 minutes washes in PBT, ovaries were incubated with a secondary antibody for 2 hours, washed 3 times x 10 minutes with PBT and lastly mounted in Vectashield with DAPI (Vector Laboratories). The following primary antibodies were used: rabbit anti-Lgl (1:100, d-300, Santa Cruz Biotechnology), rabbit anti-aPKC (1:500, c-20, Santa Cruz Biotechnology) and mouse anti-Dlg (1:200, F3 Developmental Studies Hybridoma Bank). Fixed tissue was imaged using a 1.1 numerical aperture/40x water or 1.30 numerical aperture/63x objectives on an inverted laser scanning confocal microscope Leica TCS SP5 II (Leica Microsystems) or with a 1.20 numerical aperture/63x objective on an inverted laser scanning confocal microscope Leica TCS SP8 (Leica Microsystems).

### Drug treatment

Actin cytoskeleton disruption was achieved by treating dissected ovarioles for at least 1 hour with 5 μg/mL of Latrunculin A (Sigma-Aldrich) added to the Imaging Medium (Schneider Insect Cell Medium (Gibco) supplemented with 10% FBS (ThermoFisher) and 200 μg/uL of Insulin (Sigma)). Temporal inhibition of aPKC activity in *dlg* mutant cells was achieved by using flies carrying simultaneously the *apkc^as^* allele by addition of 1NA-PP1 (Calbiochem) to the imaging media at a final concentration of 100 μM just before imaging.

### Optogenetic experiments

Flies were exposed for 24h under direct LED blue light (472 nm) prior to ovary dissection and fixation to trigger GFP-tagged protein clustering. For optogenetic depletion of PIP_2_ by the inositol polyphosphate 5-phosphatase OCRL, ovaries were dissected using a 593 nm LED light and exposed to 488 nm laser during the live imaging process. FRAP experiments were performed between around 5 minutes and 1 hour after optogenetic activation.

### Fluorescence recovery after photobleaching in the *Drosophila* follicular epithelium

*Drosophila* egg chambers from stage 7 to stage 9 were imaged in culture dishes (MatTek) with either Imaging Medium as in (Prasad et al., 2007) or with 10S Voltalef^®^ oil (VWR Chemicals). Imaging Medium was used for experiments with overexpressed GFP-tagged proteins. We resorted to the better optical properties of 10S Voltalef to improve the fluorescence signal to noise ratio in the experiments using endogenously tagged-GFP proteins. FRAP experiments were performed using a 1.1 numerical aperture/40x water objective on a Leica TCS SP5 II (Leica Microsystems) confocal microscope and using the FRAP Wizard application of the LAS Advance Fluorescence (AF) 2.6. software. The imaging plane was focused at the surface of egg chamber on the lateral cortex of the follicular epithelium (Figure S1A), where a cortical section of two neighboring cells was selected for bleaching using either a rectangular ROI with 3 μm x 1 μm size. A 1.5 μm x 1 μm ROI during the bleaching process was used to determine whether diffusion played a role in Lgl recovery (Figure 1E).

For all the FRAP experiments performed, a three-step protocol was developed: A) a prebleaching acquisition phase for GFP equilibration; B) a photoperturbation step performed using a single bleaching pulse with the 405 laser line (50 mW) at maximum laser power using the Zoom In option of the FRAP Wizard; and C) post-bleaching phases to measure the fluorescence recovery. Pre and pos-bleach imaging of GFP was performed using the 488 nm excitation line with 20%-35% laser intensity. The multi-step FRAP protocol was optimized for each basolateral polarity protein (Figure 1A), ensuring that sufficient data points were collected to accurately obtain the half-time of recovery while minimizing photobleaching during acquisition. Acquisition: Pre-bleaching – 6 frames acquired for equilibration of GFP signal (0.543s between frames); Post Bleach 1 ((1s/frame) - Lgl - 25 frames, Dlg – 50 frames; Scrib – 60 frames); Post Bleach 2 (3s/frame) – Lgl - 25 frames, Dlg – 50 frames; Scrib – 60 frames); Post Bleach 3 ((10s/frame) - Lgl - 10 frames, Dlg – 30 frames; Scrib – 35 frames).

### Image processing and fluorescence recovery curve normalization

Image processing and fluorescence intensity recovery measuring were performed using ImageJ ((Schindelin et al., 2012), (http://imagej.nih.gov/ij/)) through a pipeline composed of three main steps: 1) Registering the timelapse images. Image drift during acquisition was corrected using the “Stack Reg” plugin to align time lapse images. 2) Measuring the raw fluorescence intensity values from the corrected images. Measurements were collected in four different regions: a) 3 μm x 1 μm ROI containing the photobleached region (**Bl** in Figure S1A); b) two 3 μm x 1.5 μm ROIs in cortical regions (**NB** in Figure S1A) non-adjacent to the photobleached region to correct for photobleaching during the acquisition of post-bleaching steps; c) a 2 μm diameter circular ROI placed in the prospective nucleus, used for background subtraction as none of the proteins of interest is present in the nucleus (**B** in Figure S1A). 3) Subtracting the background, correcting for photobleaching during acquisition and normalizing recovery data. We obtain the corrected and normalized data (**%F(t)**) by applying equation (1). In this equation, background subtracted intensities in the bleached ROI **(Bl)** were normalized to the mean of the three pre-bleaching **(pre)** frames and corrected for acquisition photobleaching using the mean of the background subtracted intensities of two non-bleached ROI **(NB)**. The fluorescence recovery curve is presented as a percentage of the pre-bleach intensity, and the value of the first frame post-bleach **Bl(t_0_)** is set to zero to obtain recovery curves that enable comparison of experiments with different bleaching depths.
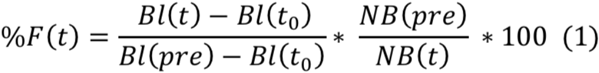

### Data analysis and statistics

The corrected and normalized data was plotted in recovery curves using GraphPad Prism 8.2.0. Recovery plots represent the average normalized fluorescence intensity and error bars indicate 95% confidence intervals (CI). We performed an extra sum-of-squares F test to assess the preferred model (bi-exponential vs single-exponential) of each individual curve obtained for Lgl-GFP fluorescence recovery. For Figure 1G, values were fitted to a <u>bi-exponentional function</u>: **F(t)= Span_Fast_ *(1-exp(-K_Fast_*t)) + Span_Slow_*(1-exp(-K_Slow_*t))**, where Span_fast_ and Span_slow_ are the amplitudes of the fast and slow fractions of recovery (constrained to a shared value between the individual curves of a single dataset) and K_slow_ and K_fast_ are rate constants expressed in reciprocal of the time. This function allows to calculate the fast half-time (t_fast_= ln(2)/K_fast_) and the slow half-time (t_slow_= ln(2)/K_slow_). As we concluded that Lgl displays a membrane-diffusion behavior, with diffusion intertwined with reactive processes (Figure 1F), we used <u>a single exponential function</u> for individual curve fitting to compare the recovery halftimes of Lgl dynamics in different experimental settings: **F(t) = Plateau * (1-exp (-Kd*t))**, where Kd represents the rate of the unbinding reaction expressed in reciprocal of the time. This function allows us to extract a simple half-time value (t^1/2^) for each curve representing one cell (t^1/2^= ln(2)/kd). We then obtained and compared half-time dispersion plots for the different biological conditions. The statistical significance between them was assessed using the non-parametric Mann-Whitney t-test.

Individual photobleaching experiments were excluded from the experimental datasets if they met one of the following criteria: 1) inefficient photobleaching step, such that reduction of intensity in the photobleached region after the bleaching pulse was below 50% (the recommended minimum for quantitative analysis of FRAP curves (McNally, 2008)); 2) low plateau of fluorescence recovery (cutoff at 65% (60% for Lgl^5A^-GFP+pmOCRL and UAS-Lgl-GFP+aPKC^ΔN^ to correct for lower plateau in the average of the whole dataset); 3) Values with low R^2^ (cutoff at 0.75 (0.65 for Lgl^5A^-GFP+pmOCRL and UAS-Lgl-GFP+aPKC^ΔN^). Mean values of the collected data are shown in Table S1. 4) the half-time of recovery is an outlier (Outliers were removed using the Extreme Studentized Method (ESD Method) with a significance level of 0.05 using a GraphPad online application (http://www.graphpad.com/quickcalcs/Grubbs1.cfm)).

## Acknowledgments

We thank Jens Januschke, Jurgen Knoblich, Stefano De Renzis, Yohanns Bellaiche, Xiaobo Wang, Daniel St Johnston and the Bloomington stock center for fly stocks. We thank Helena Richardson for detailed information on the Scrib-GFP deletion constructs, Yohanns Bellaiche for comments on the manuscript and Claudio Sunkel for assistance with supervision. This work was supported by FCT (Fundação para a Ciência e a Tecnologia) under project (PTDC/BEX-BCM/0432/2014).

## Supplementary information

Supplemental Information includes three figures, one table and three movies

